# Strong diffusional coupling between spines and dendrites promotes long-term potentiation of Schaffer collateral synapses

**DOI:** 10.1101/435446

**Authors:** Shuting Yin, Christian Schulze, Thomas G. Oertner

## Abstract

Dendritic spines on CA1 pyramidal cells are highly variable in size and shape. For some spine synapses, long and narrow spine necks provide strong diffusional and electrical isolation from the main dendrite, and it has been speculated that synapses on well isolated spines could be more plastic than synapses on spines with low resistance necks. Here we test this hypothesis by first measuring the diffusional resistance of spine necks, then pairing two-photon glutamate uncaging with bursts of back-propagating action potentials. Paired stimulation induced significant (155%) long-term potentiation (LTP) of synapses on weakly isolated spines, but no net functional change of synapses on highly isolated spines. No correlation was found between spine head volume and functional plasticity of the synapses. We conclude that contrary to our expectations, diffusional isolation makes potentiation of synapses more difficult. Our results support the concept that delivery of plasticity-related proteins from the dendrite into the spine is a limiting factor for LTP.

## Introduction

The anatomical diversity of dendritic spine shapes has been the subject of intense curiosity for a century. The concept of biochemical compartmentalization in mushroom-shaped spines has been validated in numerous functional imaging studies (1, 2). The relatively long residence time of activated enzymes in spines may keep biochemical reactions (e.g. phosphorylation events) restricted to active synapses. In recent years, evidence has accumulated that spines with long and thin necks experience strong depolarization of the head during synaptic transmission (3, 4). This strong depolarization leads to efficient unblocking of NMDA receptors and boosts calcium influx into the spine head (5–7). As high calcium concentrations are a key requirement for long-term potentiation (LTP), it is tempting to speculate that synapses on well isolated spines could have a lower threshold for LTP induction. On the other hand, successful potentiation requires an enlargement of the postsynaptic density (PSD) and insertion of additional glutamate receptors into the PSD (8, 9). Recent studies suggest that these receptors and associated scaffolding proteins are delivered from the dendrite into the spine (10, 11). It is therefore conceivable that a narrow spine neck, by restricting diffusional access to the spine, would make potentiation of the synapse more difficult (12). Whether and in which direction spine neck geometry affects the plasticity of excitatory synapses can only be determined experimentally, by coincident activation of single synapses on spines with known diffusional properties.

Spine neck dimensions are well below the resolution limit of light microscopy and thus difficult to measure in live tissue. Even state-of-the-art super-resolution approaches (13) suffer from poor resolution along the optical axis. If the measurement precision is not isotropic, the true length of the spine neck will be underestimated by a variable factor that depends on the orientation of the spine in the tissue. Furthermore, due to their variable internal organization (actin bundles, endoplasmic reticulum), even spine necks with identical outer dimensions might have different mean free path lengths for diffusing molecules (14). For our study, we did not attempt to determine the outer dimensions of spines, but instead measured the time constant of diffusional equilibration between spine head and dendrite in unperturbed neurons, before patch-clamp recordings. We compared two-photon photoactivation and fluorescence recovery after photobleaching (FRAP); both methods produced reproducible and internally consistent results.

To stimulate individual, identified synapses in a precisely timed manner, two-photon uncaging of glutamate is the method of choice. For successful LTP induction, strong Ca^2+^ influx is essential, and extracellular solutions with low or zero Mg^2+^ are usually employed to unblock NMDA receptors during plasticity induction (2, 15). This experimental trick disables the coincidence detector function of NMDA receptors. Under physiological conditions, simultaneous depolarization of the postsynaptic neuron is a necessary condition for NMDA receptor-mediated Ca^2+^ influx. Here we used artificial cerebrospinal fluid (ACSF) containing 1 mM Mg^2+^ to keep coincidence detection intact (16). We combined bursts of back-propagating action potentials (bAPs) in the postsynaptic neuron with two-photon glutamate uncaging to induce plasticity at spine synapses in CA1. The outcome of this pairing protocol was quite variable: Some synapses were significantly strengthened; others only weakly affected or even depressed. In every plasticity experiment, we also determined the diffusional resistance of the spine neck. To our surprise, synapses on highly isolated spines rarely changed their strength in response to the pairing protocol while synapses on spines with low resistance necks were much more likely to become potentiated. Thus, we can rule out the hypothesis that a high spine neck resistance promotes synaptic potentiation. Our data suggest that a long and narrow spine neck has a stabilizing effect on the weight of the resident synapse.

## Results

### Measuring diffusional coupling between CA1 dendrites and spines

CA1 pyramidal cells in organotypic slices from rat hippocampus were biolistically transfected with expression vectors encoding for the red fluorescent protein tdimer2 and photoactivatable GFP (PA-GFP, Fig. 1a). After 2-3 weeks of expression, we characterized the diffusional coupling between a number of individual spines and their parent dendrite using photoactivation of PA-GFP (810 nm) and fluorescence recovery after photobleaching (FRAP) of tdimer2 (Fig. 1b and c). The time constants were extracted by fitting a single exponential function to the time course of spine fluorescence after the laser pulse (Fig. 1d). We found that τ_decay_ of PA-GFP and τ_recovery_ of tdimer2 were highly correlated in individual spines (Fig. 1e), indicating that either measure provided us with reliable information about diffusional coupling between spine and dendrite. We used the average of τ_decay_ and τ_recovery_ as a robust measure of diffusional coupling (τ_equ_). Most spines were strongly coupled to the dendrite (small τ_equ_), highly isolated spines were quite rare. As we were interested in the functional consequences of strong diffusional isolation, we preferentially selected spines with large τ_equ_ for subsequent functional characterization and plasticity experiments.

**Figure 1.**
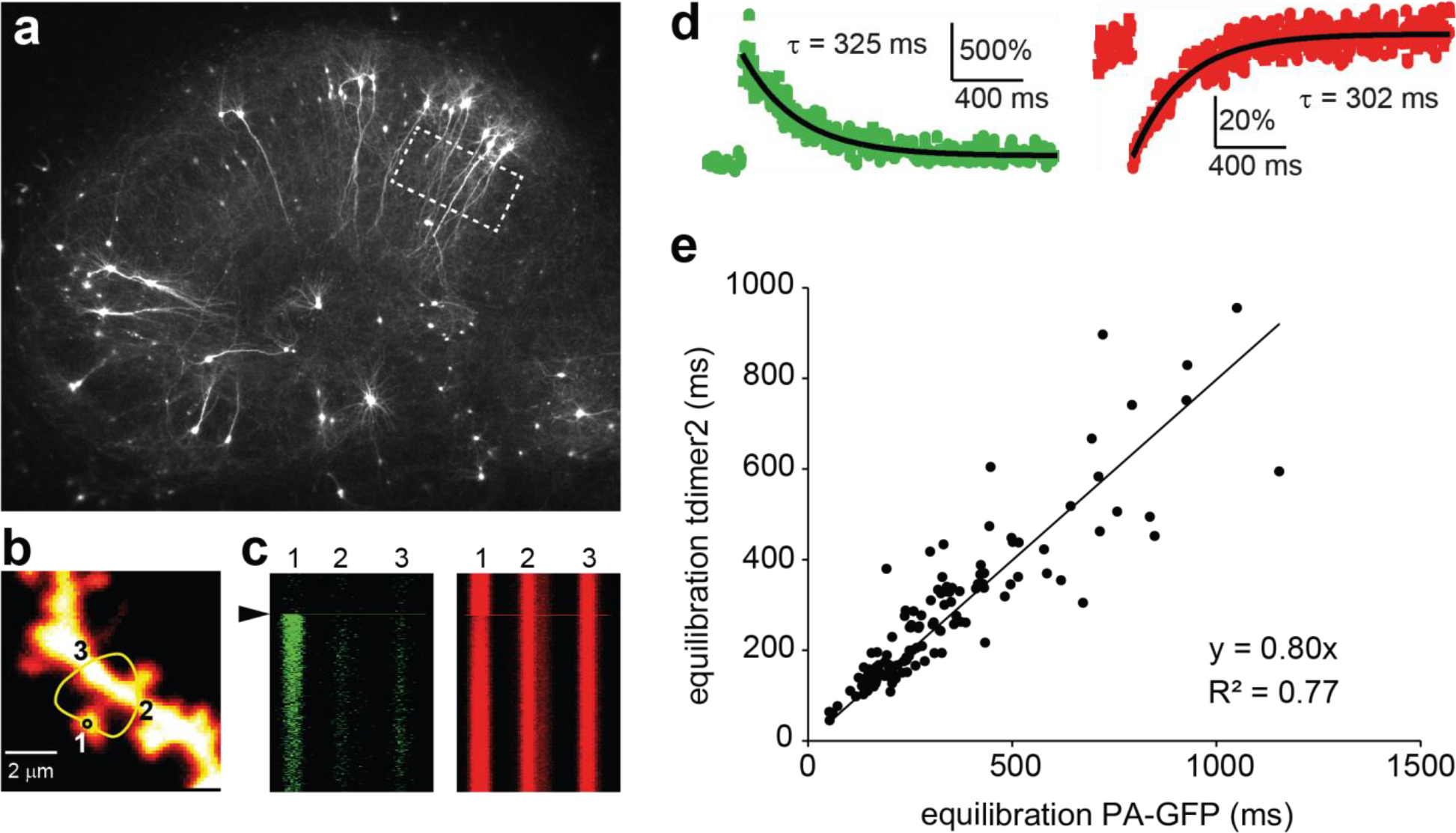
Measuring diffusional coupling between spine and parent dendrite. (**a**) Organotypic hippocampal culture after gene gun transfection with tdimer2 / PA-GFP. Box marks region of interest. (**b**) Spiny dendrite of CA1 pyramidal cell expressing tdimer2 / PAGFP at high magnification. Yellow line indicates user-defined scan path (980 nm) across spine head (1) and dendrite (2, 3). Black circle indicates position of photoactivation pulse (810 nm). (**c**) At *t* = 200 ms, a brief laser pulse (810 nm) was directed to the spine head (region 1) to simultaneously activate PA-GFP (left) and bleach tdimer2 (right). (**d**) The decay time course of PA-GFP fluorescence in the spine head (green) was very similar to the time course of fluorescence recovery after photobleaching (FRAP) of tdimer2 (red). Black lines show single exponential fits. (**e**) For individual spines, the decay time constant of PA-GFP and recovery time constant of tdimer2 were highly correlated (*n* = 120 spines, 15 cells). The distribution of τ is very broad and skewed towards strongly coupled spines (median τ_PA-GFP_ = 275 ms, range: 52 ms - 1154 ms, median τ_tdimer2_= 250 ms, range: 45 ms - 955 ms).

### Induction of LTP by pairing glutamate uncaging with postsynaptic bursts

After selecting a spine for plasticity induction, we perfused MNI-caged-L-glutamate (2.5 μM) and established whole-cell patch clamp. The red cytoplasmic fluorescence (980 nm excitation) was used to visualize dendritic morphology and to position the uncaging spot 0.5 μm from the edge of the selected spine. After break-in, we immediately started to uncage glutamate with short pulses from a second Ti:Sapph laser (720 nm, 0.5 ms). We determined the baseline potency of the synapse by collecting a series of uncaging-evoked postsynaptic currents (uEPSCs) at a rate of 1/min. After collecting baseline uEPSCs (~15 min after break-in), we paired uncaging pulses with depolarizing current injections (100 ms) to generate bursts of 7-9 back-propagating action potentials (bAPs). Paired stimulation was repeated 20 times. Synaptic potency was assessed again 30 min after the pairing protocol by a second set of uncaging pulses (Fig. 2a). Experiments were classified as significant LTP, LTD, or ‘no change’ based on the outcome of t-tests. In a first set of experiments, the pairing stimulation was repeated at 0.1 Hz. In about half of the experiments, this protocol induced a significant change in potency of the ‘paired’ synapse. A stronger protocol with 20 pairings delivered at 0.5 Hz produced an almost identical outcome (chi squared test, *p* = 0.8), suggesting that only about half of spine synapses were plastic.

**Figure 2.**
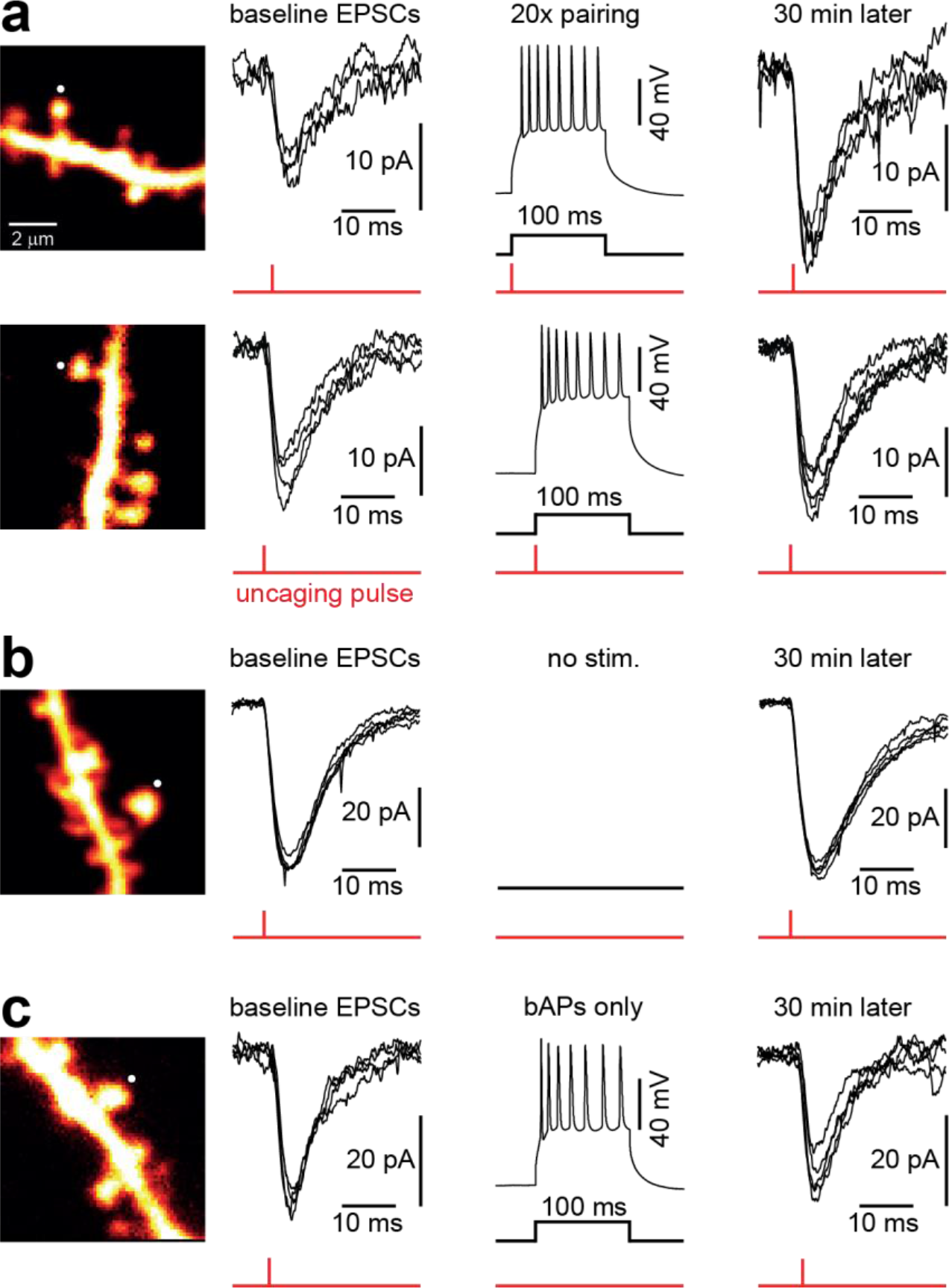
Plasticity induction by pairing of glutamate uncaging with postsynaptic depolarization. (**a**) Glutamate uncaging on a single spine (interval = 1 min) caused uEPSCs of consistent amplitude. Uncaging pulses were paired with current injections into the postsynaptic neuron (20 pairings at 0.5 Hz). Thirty minutes after the pairing protocol, uEPSCs were potentiated in some spines (upper row) whereas no plasticity occurred in other spines (lower row). (**b**) Control experiment without pairing. (**c**) Control experiment with only postsynaptic current injections, causing bursts of bAPs (20 repeats at 0.5 Hz).

As a control experiment, we assessed the stability of synaptic potency over 30 min with no stimulation (Fig. 2b). As expected, the majority of synapses (72%) maintained their strength during the 30 min (Fig. 2D), indicating that spontaneous changes in synaptic potency were rare. As an additional control, we tested whether postsynaptic spike bursts alone (20 bAP bursts at 0.5 Hz) would affect synaptic potency (Fig. 2c). There was no difference in the outcome distributions of the no stimulation and bAP groups (chi squared test, *p* = 0.2, Fig. 3a). The distribution of the paired experiments, however, was different from the control experiments (*p* = 0.0001).

**Figure 3.**
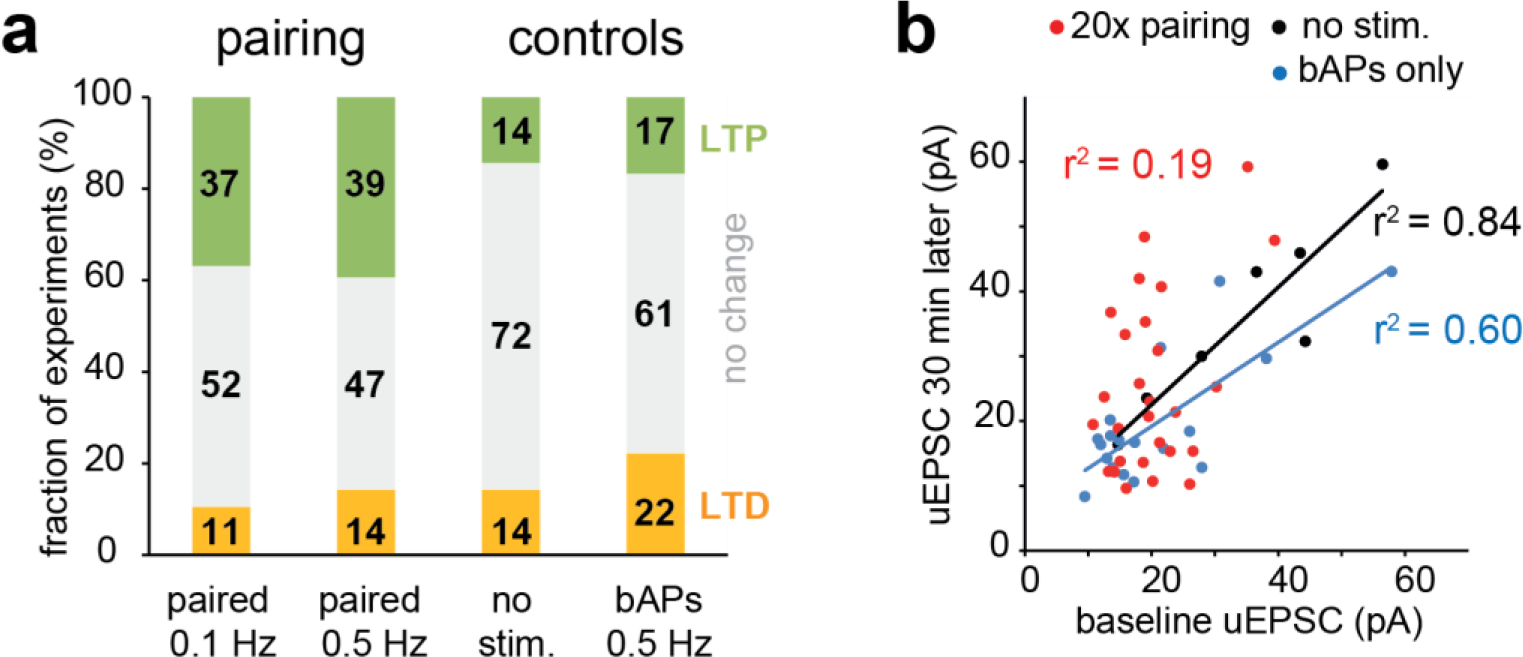
Analysis of plasticity experiments. (**a**) Outcome of uncaging-pairing stimulation and control experiments. Green indicates significant potentiation, gray indicates no change, and orange indicates significant depression. Numbers indicate % of experiments with the specified outcome (paired 0.1 Hz: *n* = 19; paired 0.5 Hz: *n* = 28; no stimulation: *n* = 7; bAPs: *n* = 18). (**b**) Under control conditions, uEPSC amplitudes at baseline and 30 min later were highly correlated (black markers, *n* = 7). Repeated action potential bursts triggered some changes, but baseline and late uEPSCs were still correlated (blue markers, *n* = 18). The pairing protocol decorrelated baseline and final potency of the synapses (red markers, *n* = 28).

Without postsynaptic stimulation, uEPSC amplitudes before and after the waiting period (30 min) were highly correlated (*r*^2^ = 0.84, Fig. 3b). Twenty trains of bAPs without simultaneous uncaging induced some plasticity, but the correlation of uEPSC amplitudes before and 30 min after bAP bursts was still high (*r*^2^ = 0.60). Paired stimulation destroyed the correlation between baseline uEPSC amplitudes and final potencies (*r*^2^ = 0.19). There was no simple relationship between initial synaptic potency and the direction or magnitude of plasticity.

### Well isolated synapses are less plastic

We were surprised by the fact that many spine synapses did not change their potency when challenged with repeated paired stimulations. We speculated that the degree of diffusional isolation from the dendrite could affect the plasticity of individual synapses. Analogous to an electrical RC circuit, the time constant of equilibration between spine and dendrite (τ_equ_) depends on the diffusional resistance of the spine neck (*R*_n_), the size of the reservoir (spine head volume, *V*), and the diffusion coefficient of the fluorescent molecule (*D*)(17):

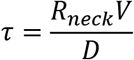

therefore

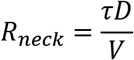

The diffusion coefficient *D* has been experimentally measured in dendritic cytoplasm (18), but it is unclear whether cytoplasmic viscosity is identical in dendrites and spines. What we know is that the diffusional resistance of the spine neck is proportional to the measured parameters *τ* and *V*:

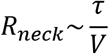

We therefore decided to report τ/*V* as a measure of spine neck resistance, in units of [s/μm^3^]. The median *R*_neck_ in our sample was 3.58 s/μm^3^, meaning it took 358 ms to exchange 63% (1/e) of the fluorescent molecules in a typical spine (0.1 μm^3^). The range was large, from 0.7 s/μm^3^ to 15 s/μm^3^, suggesting highly variable neck geometries.

Performing plasticity experiments after measuring diffusional coupling allowed us to test for correlations between the amount of plasticity induced at individual ‘paired’ synapses and the diffusional resistance of the spine neck (Fig. 4a). In a total of 28 pairing experiments, 13 synapses did not significantly change their strength (open markers, *p* > 0.05). Surprisingly, very strong potentiation (>200%) was seen in spines with relatively low neck resistance (filled markers). In the two sets of control experiments, no stimulation and bAP bursts only, spontaneous changes in synaptic strength (Fig. 4b, filled markers) were not correlated with spine neck resistance. On average, there was no net potency change in the group of unstimulated spines (106% ± 6%, *n* = 7) and in the group that was invaded by bAP bursts (102% ± 8%, *n* = 18, Fig. 4c). Therefore, we pooled these two groups (‘no stim’ and ‘bAPs only’) to form a single control group.

**Figure 4.**
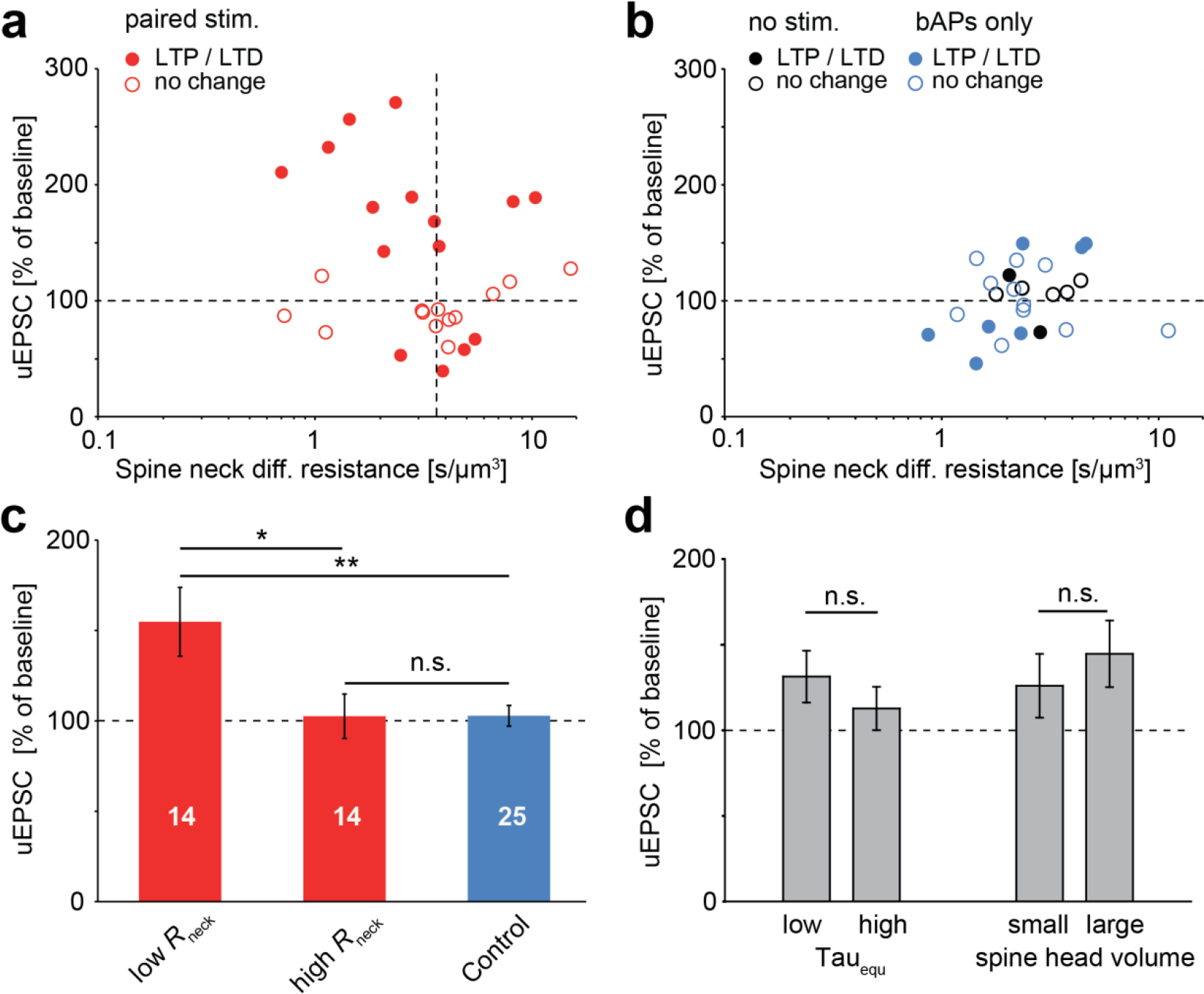
Impact of spine neck resistance on the induction of long-term plasticity. (**a**) Relationship between spine neck resistance and outcome of plasticity experiments. Filled symbols indicate experiments where uEPSC were significantly different before and after pairing at 0.5 Hz (p < 0.05), open symbols indicate experiments with no change of uEPSC amplitude. Dotted vertical line indicates median spine neck resistance of paired spines. (**b**) Unpaired control spines without stimulation (black) or twenty bursts of back-propagating action potentials at 0.5 Hz (blue). (**c**) Spines were split into two groups at the median R_neck_ (see panel A, vertical dashed line). Synapses on low R_neck_ spines were significantly potentiated compared to synapses on high R_neck_ spines (*p* < 0.05) and controls (*p* < 0.01). Synapses on high R_neck_ spines were not significantly different from the unpaired controls (from panel b). (**d**) Splitting the population of paired spines by the median time constant of equilibration (τ_equ_) or by the median volume of the spine head did not reveal significant differences between groups (low vs. high τ_equ_, *P* = 0.8; small vs. large spine heads, *P* = 0.2).

To compare the amount of plasticity induced at synapses on low and high neck resistance spines, we split the population of paired spines at the median spine neck resistance 3.58 s/μm^3^ (Fig. 4d). We found that spine synapses with low neck resistance were significantly potentiated compared to controls (155% ± 19%, *p* = 0.01, ANOVA followed by t-tests), while spine synapses with high neck resistance were not (103% ± 12%, *p* = 0.99).

As we calculated spine neck diffusional resistance from the time constant of equilibration (τ_equ_) and the spine head volume, we asked whether one of these directly measured parameters could already explain the differences in plasticity. However, splitting the population of paired spines at the median τ_equ_ did not reveal a significant difference in plasticity between the two groups (Fig. 4e, *p* = 0.8), and neither did spine head volume (*p* = 0.2). Thus, spine neck resistance was the only structural parameter correlated with synaptic plasticity, albeit not in the direction we expected.

We were concerned that the differences in plasticity might be due to other factors than *R*_neck_. Did we stimulate one set of spines more strongly? The laser pulse used for glutamate uncaging bleaches some of the red fluorescent protein in the spine. The degree of bleaching, which is a measure of local two-photon activation and therefore proportional to the amount of glutamate released, was not different between the two groups (low *R*_neck_ group: 23% ± 4% bleach, high *R*_neck_ group: 25% ± 4% bleach, *p* = 0.7), indicating stimulation strength was not systematically different. Was wash-out different in the two groups? The time between break-in and start of the induction protocol was not different between groups (low *R*_neck_ group: 17 ± 2 min, high *R*_neck_ group: 15 ± 1 min, *p* = 0.4), excluding wash-out as a possible explanation for the observed difference in plasticity. Is it possible that synapses on high *R*_neck_ spines were stronger to begin with and therefore more difficult to potentiate? There was no difference in initial potency (low *R*_neck_ group: 19.6 ± 2.1 pA, high *R*_neck_ group: 20.4 ± 1.3 pA, *p* = 0.7). Spine head volumes, however, were significantly larger in low *R*_neck_ spines (low *R*_neck_: 0.18 ± 0.03 μm^3^, high *R*_neck_ group: 0.08 ± 0.01 μm^3^, *p* = 0.003). Consistent with this finding, a positive correlation between spine head volume and spine neck diameter was described in earlier two-photon studies (19) and in a recent serial EM study of perfusion-fixed hippocampal tissue (20). Thus, even though spine neck geometry is not completely independent of spine head size, we found that only *R*_neck_ had a measurable impact on synaptic plasticity.

## Discussion

Our pairing protocol had overall a relatively modest effect on synaptic strength. To maintain coincidence detection by NMDA receptors, we used ACSF containing 1 mM Mg^2+^ and paired uncaging pulses with brief bursts of postsynaptic action potentials. Under these conditions, calcium influx is boosted and plasticity is triggered if the first back-propagating AP arrives within 40-50 ms after the EPSP (6, 23). Until now, many studies have used glutamate uncaging in magnesium-free solution (2, 15, 21, 22). This protocol leads to strong Ca^2+^ influx and LTP at every stimulated spine. However, the potential influence of the spine neck on spine head depolarization cannot be investigated with uncaging in zero Mg^2+^ since this circumvents the requirement for coincidence detection by NMDA receptors. In physiological Mg^2+^, the calcium rise induced by the interaction of back-propagating APs and glutamate receptors is supra-linear and most pronounced in highly isolated spines. We therefore expected that spines with high *R*_neck_ would be more likely to undergo LTP and were surprised by the lack of positive correlation. It should be noted that we actively biased our sample towards highly isolated spines, which are relatively rare across the population. A ‘blind’ electrophysiological LTP experiment would therefore be dominated by synapses on low *R*_neck_ spines, which did produce substantial (155%) potentiation in our experiments. We performed somatic voltage-clamp recordings to measure uncaging-evoked synaptic currents, but acknowledge the fact that spine heads escape the voltage clamp when responding to glutamate (5, 24). Since we were interested in changes before and after pairing rather than determining the absolute number of AMPA receptors in a synapse, we considered it important to keep the dendritic membrane potential as constant as possible over the entire experiment (~45 min). A consequence of escaping (i.e. depolarizing) spine heads is that the measured LTP may underestimate the true increase in synaptic conductance.

Our results challenge the widely held belief that spine necks have evolved to enhance synaptic plasticity (19). The high input resistance and biochemical isolation, it was argued, would trap calcium-activated enzymes such as CaMKII close to their targets at the synapse, facilitating LTP. Our data suggest this trapping effect is not sufficient to enable LTP at the most isolated synapses. Rather the opposite seems true, and less isolated spines are more likely to increase their AMPA currents after pairing. The major delivery route for spine AMPA receptors is lateral diffusion in the membrane rather than constitutive exocytosis in the spine head itself. The spine neck restricts diffusion of receptors and other membrane proteins into and out of mushroom spines (12, 25). Restricted access to AMPA receptors and other plasticity-related proteins is a plausible explanation for our finding that synapses on highly isolated spines were less plastic.

## Materials and Methods

### Organotypic hippocampal cultures

Hippocampal slice cultures were prepared in accordance with the ethical standards of the German Animal Welfare Act and the State Authority of Hamburg, Germany, using female rat pups (Wistar) at postnatal day 5-6. The hippocampus was dissected as described (28) and cut in 400 μm thick slices using a McIlwain tissue chopper. No antibiotics were used during the preparation or in the culture medium. At DIV 7-9, cultures were transfected with a 1:1 mixture of expression vectors encoding tdimer2 (29) and PA-GFP (30), each driven by the synapsin-1 promoter, using a Helios gene gun (Bio Rad).

### Two-photon imaging and photoactivation

After 2-3 weeks of expression (DIV 24-30), photoactivation experiments were performed on a custom-built two-photon microscope controlled by ScanImage 3.8 open source software (31). We extended the capabilities of ScanImage to perform user-defined curved line scans and automatic drift correction. Two Ti:Sapph lasers (MaiTai DeepSee, Spectra-Physics) were coupled into the microscope through a polarizing beamsplitter cube (Thorlabs), controlled by electro-optical modulators (Conoptics) for simultaneous imaging (980 nm), photoactivation (810 nm) or uncaging (720 nm) of MNI-caged-L-glutamate (Invitrogen). Photons were collected through objective (60x 0.9 NA, Olympus) and condenser (1.4 NA, Olympus) and detected with 4 GaAsP PMTs (Hamamatsu). During imaging and plasticity induction, cultures were submerged in artificial cerebrospinal fluid (ACSF) containing (in mM): 127 NaCl, 2.5 KCl, 2 CaCl_2_, 1 MgCl_2_, 25 NaHCO_3_, 1.25 NaH_2_PO_4_, 25 D-glucose, 2.5 MNI-caged-L-glutamate (pH 7.4, ~308 mOsm, saturated with 95% O_2_ / 5% CO_2_).

### Spine head volume measurements

We imaged spines in *stratum radiatum*, located on thin oblique dendrites of CA1 pyramidal cells. We used the integrated fluorescence of the spine head as a measure of spine volume. Spine fluorescence (*F*_spine_) was determined from a stack of images and normalized by the maximum fluorescence from the same cell (*F*_max_), obtained by immersing the point spread function (PSF) in the trunk of the apical dendrite. The volume of the PSF (*V*_PSF_ = 0.18 fl) was determined from images of 0.1 μm fluorescent microspheres (Molecular Probes) excited at 980 nm. The volume of the spine head (*V*_spine_) was calculated as

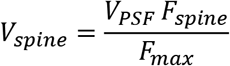

### Electrophysiology and pairing protocol

Patch-clamp recordings were performed at 32°C under Dodt contrast using a Multiclamp 700B amplifier (Axon Instruments) and Ephus open source software (32). Patch pipettes were filled with pipette solution consisting of (in mM): 135 K-gluconate, 4 MgCl_2_, 4 Na_2_-ATP, 0.4 Na-GTP, 10 Na_2_-phosphocreatine, 3 ascorbate, and 10 HEPES (pH 7.2, 295 mOsm). The power of the 720 nm uncaging laser was adjusted to bleach ~25% of red spine fluorescence during a 1 ms pulse directed to the outer edge of a test spine. The test spine was not used for physiological experiments. LTP was induced by pairing of 0.5 ms uncaging pulses with 100 ms depolarizing current injections (20 repeats at 0.1 Hz or 0.5 Hz). The potency of the synapse was assessed 30 min after the pairing protocol (5 test pulses, 1/min). Input and access resistance were continuously monitored; experiments with >50% change of either value were excluded from further analysis. As a control group, we assessed the stability of synaptic responses on spines that were invaded by bAPs, but not simultaneously activated by glutamate uncaging. Some of these spines were on neurons where no spine received paired stimulation (*n* = 10), others on the same dendrite as a paired spine, but at >10 μm distance (*n* = 8).

### Data analysis

Analysis of line scan data and curve fitting was performed with custom software written in Matlab. We used GraphPad Prism for statistical analysis. Long-term plasticity of single synapses was assessed by comparing uEPSP amplitudes at baseline to uEPSP amplitudes 30 min after pairing (Student’s *t*-test, two-tailed). On the population level, significance was tested by Chi squared tests or by ANOVA followed by two-tailed Student’s *t*-tests. We considered *p* < 0.05 as significant (*) and *p* < 0.01 as highly significant (**). Data are shown as mean ± standard error (SEM).

## Author contributions

S.Y. performed experiments and analyzed data, C.S. developed software for the laser scanning microscope, T.G.O. designed the study and wrote the manuscript.

## Data Availability

The authors will make available all data and materials. Interested parties should contact T.G.O. to make requests.

## Acknowledgements

We thank Dr. Christine E. Gee for critical reading of the manuscript and Iris Ohmert for excellent technical support. This study was supported by the German Research Foundation (DFG) through Priority Programme SPP 1665, Collaborative Research Center SFB 936, Research Unit FOR 2419, and by the Ministry of Science, Research and Equalities Hamburg (Z-FR LF).

